# Comparative transcriptome profiles of first-stage larvae and adult female *Dracunculus medinensis* (Guinea worm)

**DOI:** 10.1101/2025.08.01.668080

**Authors:** Hassan Hakimi, Sabina Beilstein, Michael J. Yabsley, Christopher A. Cleveland, Lucienne Tritten, Guilherme G. Verocai

**Author notes:** Corresponding authors: Lucienne Tritten, PhD, Guilherme G. Verocai DVM, MSc, PhD, DACVM. These authors contributed equally to this study (Hassan Hakimi, Sabina Beilstein).

## Abstract

*Dracunculus medinensis*, also called the Guinea worm, is a nematode that causes dracunculiasis, a debilitating neglected tropical disease in humans. The parasite is currently targeted by the global Guinea Worm Eradication Program (GWEP). Historically, GWEP in endemic countries have focused on interrupting transmission of the disease through intervention such as isolation and management of patients, health education, provision of improved water sources and promotion of filtering drinking water to avoid ingestion of the copepod intermediate host (IH) that may contain infectious third-stage larvae. The recent shift of Guinea worm infections in animals - particularly domestic dogs - has introduced an additional challenge to the eradication program, underscoring the urgent need for diagnostics and therapeutics. Understanding the parasite biology and survival strategies in the mammalian host, the copepod IH, and fresh water is pivotal to identifying new control measures. Comparative transcriptomic analysis provides a powerful tool to uncover the molecular mechanisms underlying parasite survival and adaptations. Here, we compared the transcriptome of adult gravid female and first-stage larvae (L1), the stage infective for the copepod IH. Comparative transcriptomic analysis of two adult females and their L1 revealed an upregulation of genes involved in translation, transcription, and DNA repair in L1, likely reflecting adaptations essential for survival in freshwater and subsequent infection of copepods. Additionally, genes involved in cuticle formation were upregulated in adult females highlighting the role of cuticle integrity in retaining millions of L1 until the gravid female worm emerges. We identified highly expressed genes in the adult female that may represent promising candidates for diagnostic markers. This study provides novel insights into the biology of the Guinea worm by examining the transcriptome of L1 and adult female stages. These findings could support the development of novel diagnostics and therapeutics to advance the ongoing eradication effort.

**Authors summary:** Guinea worm disease is caused by the nematode *Dracunculus medinensis,* a parasitic worm responsible for a debilitating neglected tropical disease in humans and targeted for global eradication. Infection occurs through the consumption of contaminated drinking water with the infective larval stage harbored within fresh water crustacean copepods - the intermediate host of the parasite. The high number of Guinea worm infections in animals specially dogs poses a significant challenge to eradication, as infected animals act as reservoirs, contributing to the parasite’s persistence in the environment. Furthermore, the absence of early diagnostic tools and effective therapeutics complicates disease control. In this study, we performed a comparative transcriptome analysis of adult female Guinea worms and their first-stage larvae. We identified highly expressed genes in the adult female that may represent promising candidates for diagnostic markers. Additionally, we found genes and pathways upregulated in first-stage larvae which are likely essential for survival in freshwater and subsequent infection of copepods. Our study provides insight into the molecular mechanisms underlying Guinea worm survival across life stages and environments. These findings aim to support the development of novel diagnostics and therapeutics to advance ongoing eradication efforts of this neglected tropical disease.

## Introduction

Dracunculiasis or Guinea worm (GW) disease is a painful and debilitating neglected tropical disease of humans caused by the nematode *Dracunculus medinensis*. GW belongs to the superfamily Dracunculoidea in the order Spirurida in Clade III of Nematoda which includes important pathogenic human filarioid nematodes, *Onchocerca volvulus*, *Wuchereria bancrofti*, and *Brugia malayi* [1]. The global GW Eradication Program (GWEP), launched in the 1980s, has successfully reduced human cases of the disease over the past four decades from an estimated 3.5 million cases in 1986 to just 15 cases in 2024 [2]. In the absence of an effective drug or vaccine, the eradication program has relied on case containment and preventive measures, including health education to promote safe drinking practices and the treatment of water sources with the organophosphate larvicide temephos (Abate^®^) to decrease copepod populations [3–5].

GW has a complex lifecycle involving a mammalian definitive host (DH) and a cyclopoid copepod intermediate host (IH). The gravid adult GW female can reach up to one meter in length within subcutaneous connective tissues of their DH, from which it emerges through the opening of a blister to release first-stage larvae (L1) into water. For further development, these L1 must be ingested by small freshwater crustacean copepods (Copepoda; Cyclopidae) [6]. The development of L1 to infective third-stage larvae (L3) within the copepod IH takes ∼two weeks under optimal environmental conditions [5, 6]. Upon ingestion of L3-infected copepods by a DH, the larvae are released in the stomach, penetrate the intestinal wall, and migrate through the peritoneal cavity, where they mature into adults. Following mating, the adult female worm develops and migrates along muscle planes, often to the legs, emerging through the skin 10-14 months after infection [6, 7].

GW was long considered an exclusively human parasite with no animal reservoir [7], although experimental infection of dogs was successful [8–10]. Historically, natural infections in dogs were also reported in endemic regions yet these cases were reported to have disappeared following elimination of human infections in these specific areas [8–10]. However, active village-based surveillance in Chad revealed 27 dog infections in 2012 primarily in outbreaks along the Chari River, with the number of dog cases increasing to 1,480 in 2020 [8, 11]. Genomic sequencing of nematodes infecting both dogs and humans confirmed that these belong to the same species, and that the parasite circulates between humans and dogs [12]. Additionally, other endemic countries reported cases in domestic dogs and cats, as well as wild baboons [10, 13]. In addition to the ingestion of copepods through drinking water, the involvement of paratenic or transport hosts - such as amphibians and fish – likely contributes to the unusual epidemiology of GW in dogs around the Chari River [8, 14–16]. Despite the low incidence in humans, the number of animal cases remains high, with 662 animal infections reported from endemic countries mainly from Chad and Cameroon in 2024 [2]. The circulation of GW in animal populations poses a risk of human exposure and hampers eradication efforts.

Although the GWEP was successful in significantly reducing the number of human cases, challenges remain in certifying the eradication of GW in the five remaining endemic countries (Angola, Chad, Ethiopia, Mali, and South Sudan) in Africa. The persistent occurrence of GW in animal population, particularly domestic dogs, the absence of diagnostic tools to detect infections before the emergence of the adult female worm, and lack of effective anthelmintic drugs for treating infected dogs remain key obstacles that must be addressed through a comprehensive One Health approach. These emerging challenges have been acknowledged by the GWEP through a reevaluation of its research agenda [17].

GW circulates between a warm-blooded mammalian host and a freshwater copepod as an intermediate host. Changes in gene expression make this transition possible. Insights into GW survival in both the mammalian DH and copepod IH are crucial for advancing our understanding of the parasite and guiding the development of targeted control strategies. The genome of an adult female GW from the Republic of Ghana was sequenced by the International Helminth Genomes Consortium with 103 Mbp encoding for ∼ 11,000 proteins [18]. Additional GW genome sequences from other endemic countries including Chad, are also available [12].

In the present study, we used next-generation sequencing to comparatively analyze the transcriptomes of L1 and adult female stages of GW. We identified stage-specific pathways that were upregulated in the L1 stage, which are likely required for survival in the freshwater environment and subsequent acquisition by the copepod IH. Additionally, we found genes involved in cuticle formation upregulated in the adult female, likely playing a role in maintaining the cuticular integrity and retaining L1 until emergence into the environment. This data set provides a foundation for future studies to identify novel biomarkers and therapeutic targets.

## Methods

### Collection of parasite samples and RNA extraction

A ferret (*Mustela furo*) was experimentally infected at the University of Georgia (IACUC number: A2020 05-003) by feeding copepods infected with L3 stage of *D. medinensis* [14, 19]. Two subcutaneous female worms were recovered from the exposed ferret upon necropsy. Five pools of L1 stage were prepared from the two adult worms, counted, and stored at -80°C pending RNA extraction. Samples were thawed on ice and adult female tissues and L1 pools were homogenized with liquid N2 flash freezing and manual grinding using disposable pestles. Total RNA was extracted using the Total RNA Purification Kit (Norgen Biotek Corp., Thorold, ON, Canada) according to the manufacturer’s instructions. RNA was quantified using Quantus^TM^ Fluorometer (Promega, Madison, WI) and stored at -80°C until being shipped on dry ice to LC Sciences (LC Sciences, Houston, TX) for mRNA sequencing. mRNA sequencing was performed by LC Sciences Poly (A) RNA-seq Sequencing Service.

### Library preparation and RNA sequencing

Poly(A) RNA sequencing library was constructed following Illumina’s TruSeq-stranded-mRNA sample preparation protocol. RNA integrity was checked using Agilent Technologies 2100 Bioanalyzer. Poly(A) tail-containing mRNA was purified using oligo- (dT) magnetic beads and fragmented using divalent cation buffer in elevated temperature. It was followed by cDNA synthesis, second strand synthesis, and adaptor ligation. The ligated products were amplified by PCR. Quality control analysis and quantification of the sequencing library were performed using Agilent Technologies 2100 Bioanalyzer High Sensitivity DNA Chip. Paired-end sequencing was performed on Illumina NovaSeq 6000 sequencing system.

### Transcript assembly

Adapter trimming and removal of undetermined or low-quality bases were performed using Cutadapt [20] and an LC Science in house perl script. FastQC (http://www.bioinformatics.babraham.ac.uk/projects/fastqc/) was used for sequence quality control. Reads were mapped to the reference genome of *Dracunculus medinensis* from WormBase [21] using HISAT2 [22]. Mapped read assembly was performed using StringTie [23], followed by the merging of all transcriptomes to reconstruct a comprehensive transcriptome using Perl scripts and *gffcompare* [24]. To estimate the expression levels of all transcripts, StringTie [23] and ballgown (http://www.bioconductor.org/packages/release/bioc/html/ballgown.html) were used.

### Differential gene expression analysis

mRNA expression levels were analyzed using StringTie [23], with quantification based on FPKM values. Differential expression analysis was performed with the R package DESeq2 [25] between two groups and with the R package edgeR [26] between two samples. mRNAs were considered differentially expressed if the false discovery rate (FDR) was < 0.05 and the absolute fold change was ≥ |2|.

### Gene expression analysis using ExpressAnalyst

For gene expression analysis we used the online platform of ExpressAnalyst [27–30] designed to perform comprehensive gene expression and meta-analysis. Unprocessed data of adult and larval L1 GW were used as input. The ExpressAnalyst platform relies on the Seq2Fun algorithm (version 2.0.5) [29, 30], which is a functional profiling tool for RNA-seq data analysis for organisms without reference genome. For *D. medinensis,* we must rely on a poorly annotated genome. Seq2Fun relies on pre-built databases. The nematode group, which we used, contains 6 species including *Caenorhabditis elegans*, *Caenorhabditis briggsae*, *B. malayi*, *Loa loa*, *Necator americanus*, and *Trichinella spiralis* (https://www.seq2fun.ca/database.xhtml, accessed November 2024).

ExpressAnalyst generates a counts table with Seq2Fsun ortholog IDs as output following raw data processing. This expression table was further used to proceed to a generic and species independent statistical and functional analysis in the ExpressAnalyst environment. Default values were used for filtering out features with an abundance threshold set to 5 counts or lower and a variance percentile rank at 15 or lower. For basic whole transcriptome expression profiling, normalization was performed using Trimmed Mean of M-values (TMM) method to enable PCA visualization. Differential gene expression analysis was done using the built-in EdgeR version [26] with preset parameters (ExpressAnalyst). Pairwise comparison was conducted in adult GW female against L1. A log_2_ fold-change threshold of 0 was applied (including all hits with a log_2_ fold-change greater than 1), along with an adjusted P-value cut-off of 0.05.

At the functional level, pathway analysis and visualization were further conducted within the ExpressAnalyst environment using GSEA (Gene Set Enrichment Analysis) with the GO:BP (Gene Ontology Biological Process) and GO:MF (Gene Ontology Molecular Function) databases.

### Enrichment Analysis

The output file from ExpressAnalyst containing the differentially expressed genes was used for further enrichment analysis of the mRNA sequencing data using EnrichR (version enrichR_3.2) in R Studio (version 4.4.1). Differentially expressed genes were separated into enriched and depleted according to their log2 fold-change. We selected four databases: GO:BP, GO:MF, MSeqDB Hallmark (Molecular Signatures Database), and Reactome. Visualization was done using ggplot2 (version 3.5.1) in R Studio (version 4.4.1).

### Identification of novel diagnostic marker candidates

To identify potential diagnostic markers, genes aligned to the GW reference Genome were ranked based on their expression levels in the adult female stage (S1 Table). These genes were then screened for putative signal peptide using SignalP 5.0 [31] and transmembrane domains were predicted using TMHMM-2.0 [32].

## Results

### Transcriptome profile of *D. medinensis*

Over 440 million reads were generated from total RNA extracted from *D. medinensis* L1 and adult female stages isolated from a laboratory ferret infection. Following removal of low-quality reads and reads that were mapped on the ferret genome, 84.6% of sequenced reads mapped to *D. medinensis* genome (Table 1). Of 10,970 predicted genes in the *D. medinensis* genome, 69.6-81.7% of genes were expressed in L1 or adult female stage. The median gene expression in L1 was 80.6% which was higher than in adult female with a median expression of 70.2%.

**Table 1.** Total number of reads sequenced and mapped to the *D. medinensis* genome.

| Sample* | Million reads sequenced | Million reads mapped | Expressed genes | Mean coverage (FPKM)/Transcript | *D. medinensis* transcripts detected (%) |
| --- | --- | --- | --- | --- | --- |
| GW1-L1-1 | 40.1 | 34.0 (84.77%) | 8961 | 158.19 | 81.7 |
| GW1-L1-2 | 40.2 | 33.7 (83.81%) | 8843 | 158.15 | 80.6 |
| GW2-L1-1 | 39.6 | 33.2 (83.77%) | 8852 | 156.83 | 80.7 |
| GW2-L1-2 | 41.3 | 36.1 (87.46%) | 8428 | 167.59 | 76.8 |
| GW2-L1-3 | 39.7 | 33.7 (84.87%) | 8576 | 171.35 | 78.2 |
| GW1-1 | 39.8 | 34.1 (85.57%) | 7642 | 156.79 | 69.7 |
| GW1-2 | 39.9 | 33.7 (84.36%) | 7694 | 160.17 | 70.1 |
| GW1-3 | 40.6 | 34.1 (83.89%) | 7697 | 157.91 | 70.2 |
| GW2-1 | 39.5 | 32.6 (82.54%) | 7769 | 155.51 | 70.8 |
| GW2-2 | 40.7 | 34.9 (85.81%) | 7636 | 160.13 | 69.6 |
| GW2-3 | 39.0 | 32.6 (83.44%) | 7892 | 153.96 | 71.9 |

\* GW1 and GW2 are two individual worm retrieved from a ferret experimental infection with three technical replicated. Five pools of L1 in total were collected from GW1 and GW2.

### Stage-dependent differential gene expression

Given that the currently available GW genome is not well annotated, gene expression analysis was done using ExpressAnalyst [27–30] by comparing L1 and adult female GW. Principal component analysis (PCA) showed that the stage accounted for most of the variation in datapoints. The L1 originating from two distinct GW females showed very close proximity and it seems that the maternal origin accounts for less variation in L1 stages compared to the adult form which shows less proximity (Fig 1).

**Fig 1.**
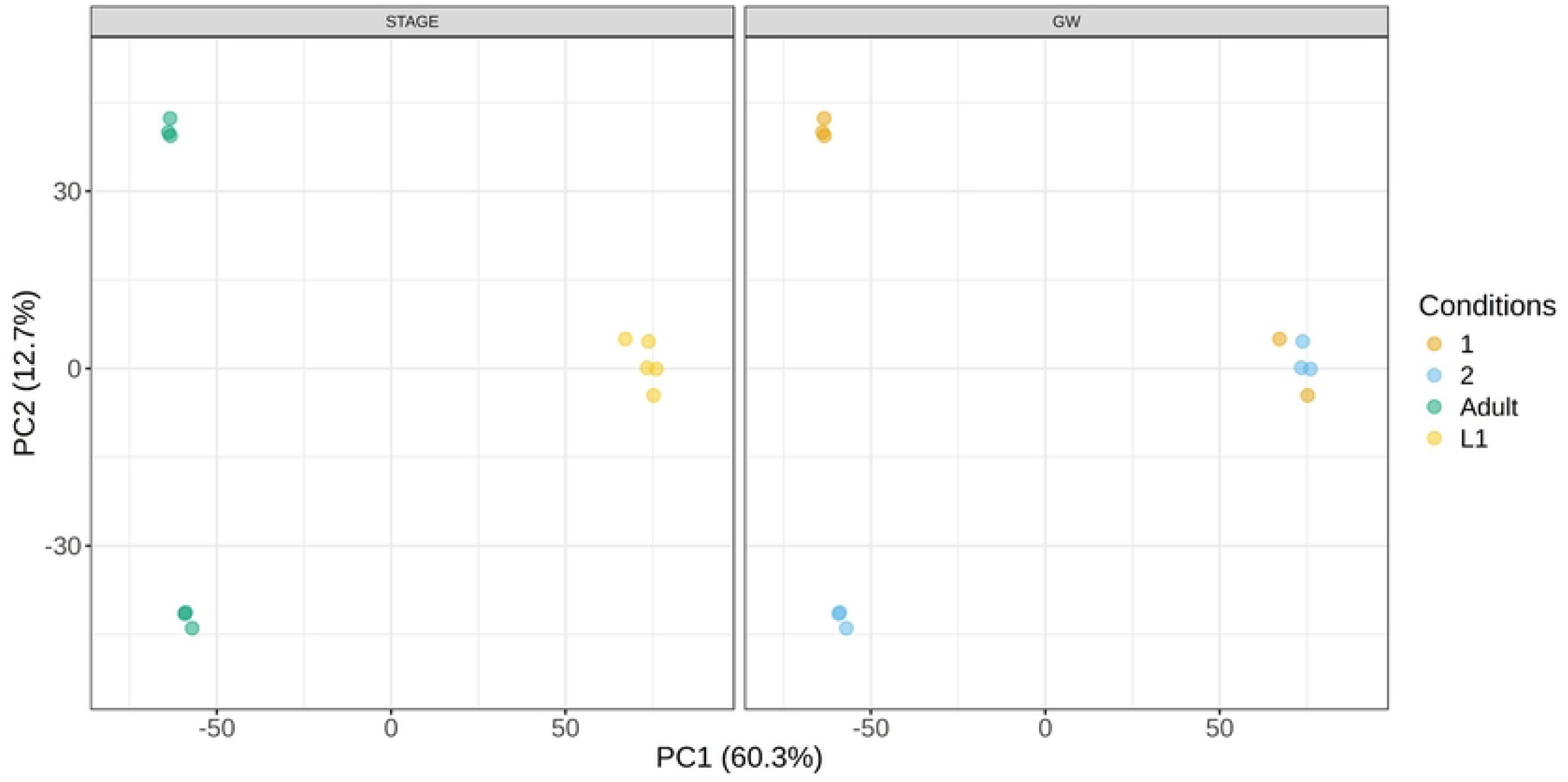
Principal component analysis (PCA) of adult female and L1 stages transcriptome. PCA is shown after submitting the gene expression data to Trimmed Mean of M-values (TMM) normalization in ExpressAnalyst. The PCA plot was created using the ExpressAnalyst platform (Seq2FunM 2.0.5). STAGE, Adult (specimen 1 and 2) and L1 stages of *D. medinensis*; GW, Guinea Worm specimen 1 and 2.

Differential gene expression analysis done using ExpressAnalyst on a total of 5454 features identified based on the ortholog mapping revealed 4339 significant features. The number goes down to 2457 with a log_2_ fold change set to 0 and a significance level of P_adj_ < 0.05 (S1 Fig).

Gene set enrichment analysis (GSEA) using ExpressAnalyst revealed differentially expressed pathways when accessing the two different databases, GO:BP (Gene Ontology Biological Process) and GO:MF (Gene Ontology Molecular Function) (Table 2) [33, 34]. For GO:BP only two pathways showed up, neuropeptide signaling pathway and translation. In contrast, GO:MF showed five different pathways: structural constituent of cuticle, DNA binding transcription factor activity specified as RNA polymerase II specific, structural constituent of ribosomes, sequence specific DNA binding, and helicase activity. The numbers of GO terms appearing from our data analysis range from 13 to 153 hits corresponding to the specific pathway with their different total numbers of GO terms assigned to the pathway. Six out of seven hits showed a negative enrichment score over the two analyses indicating downregulation of genes in those pathways in adults compared to L1 larval stages. Heatmaps generated show the genes contributing to the pathway analysis (S2 Fig).

**Table 2.** Gene Set Enrichment Analysis (GSEA) using two databases.

| **Database: GO Biological Process** |  |  |  |  |  |
| --- | --- | --- | --- | --- | --- |
| Name | Total | Hits | EnrichmentScore | Pval | Padj |
| 1 Neuropeptide signaling pathway | 287 | 14 | -0.821 | 0.000203 | 0.01591 |
| 2 Translation | 939 | 153 | -0.402 | 0.000441 | 0.02392 |

| **Database: GO Molecular Function** |  |  |  |  |  |
| --- | --- | --- | --- | --- | --- |
| Name | Total | Hits | EnrichmentScore | Pval | Padj |
| 1 Structural constituent of cuticle | 400 | 13 | 0.826 | 0.000198 | 0.00631 |
| 2 DNA_binding transcription factor activity, RNA polymerase II_specific | 1738 | 47 | -0.694 | 0.000208 | 0.00631 |
| 3 Structural constituent of ribosome | 751 | 96 | -0.492 | 0.000215 | 0.00631 |
| 4 Sequence_specific DNA binding | 1271 | 40 | -0.554 | 0.000834 | 0.01843 |
| 5 Helicase activity | 621 | 67 | -0.467 | 0.001067 | 0.02124 |

Pval, P-value; Padj, adjusted P-value. Analysis was done with ExpressAnalyst (Seq2FunM 2.0.5).

### Functional characterization of differentially expressed genes and pathways

Using the output data from ExpressAnalyst, containing a list of genes, an enrichment analysis was done using EnrichR [35–37]. The analysis incorporated a broader range of databases compared to the GSEA analysis performed using ExpressAnalyst. MSigDB Hallmark [38–40] and Reactome [41, 42] were additionally included in this analysis. Enriched and depleted significant features in the adult compared to the L1 stage were analyzed separately. The 10 most enriched or depleted pathways were considered in the graphical representation (Fig 2). The results varied in adjusted P-values across the different databases, with the Reactome database yielding the most significant findings for both depleted and enriched pathways (S3 Fig). In contrast, the MSigDB Hallmark database showed lower significance in both datasets compared to the results from the other databases. Among the top enriched features, two data points from GO:BP stand out, as they both show a high gene ratio contributing to the pathway, namely the Oligosaccharyl Transferase Activity and the NEDD8 Transferase Activity. A similar pattern was observed for Ubiquitin Ligase Inhibitor Activity and the Ubiquitin-Protein Transferase Activity, which emerged in the GO:BP analysis of the depleted features. The pathway Establishment of Protein Localization to Endoplasmic Reticulum also exhibits a relatively high gene ratio compared to other enriched pathways in the GO:MF database. The same is true for Cytoplasmic Translation among depleted biological features analyzed using the GO:MF database. In addition, it is also the most significant hit in the depleted dataset.

**Fig 2.**
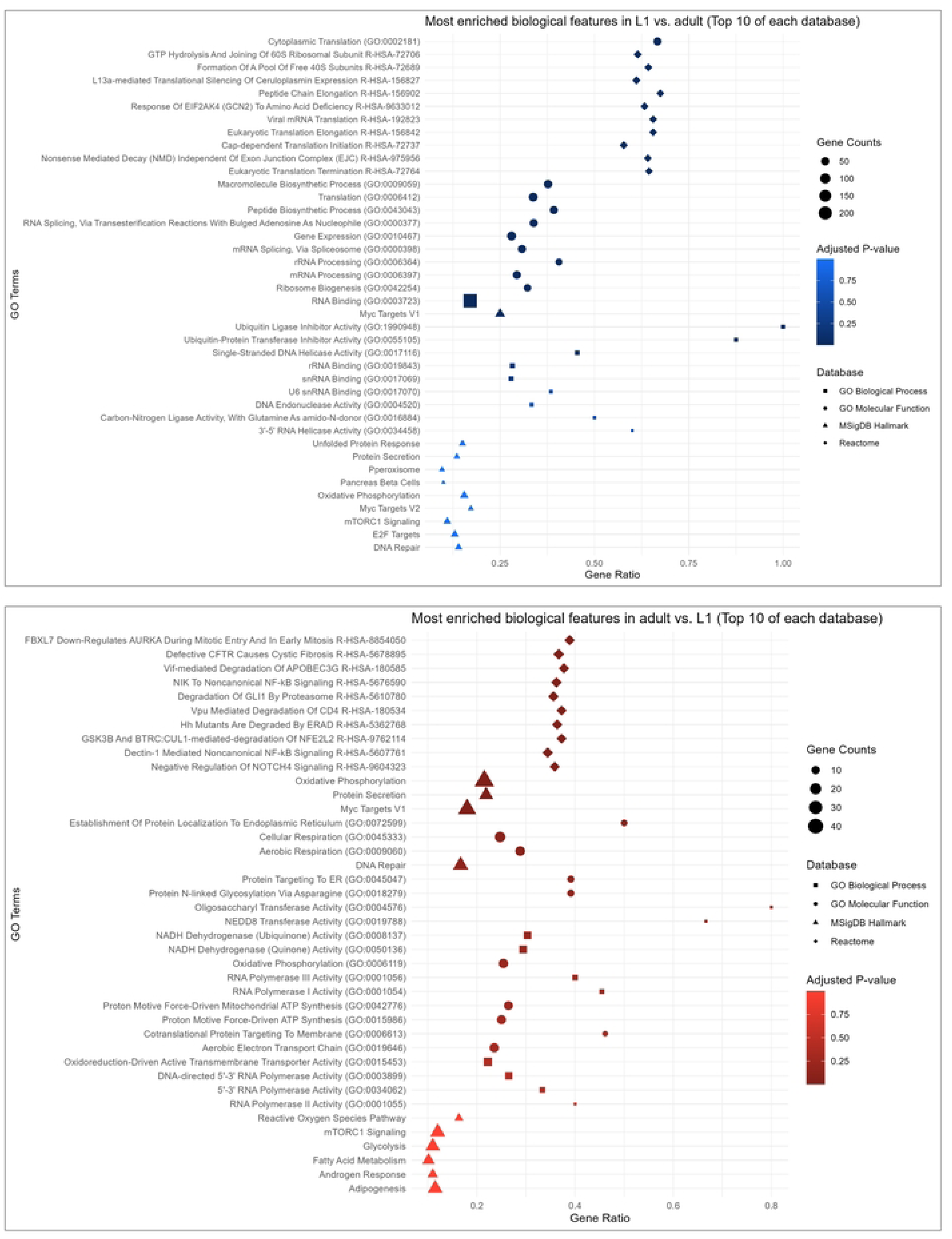
Dot plot of enriched biological pathways in adult or L1 stages. Pathway enrichment analysis of differentially expressed genes in adult compared to larval L1 stages of *Dracunculus medinensis*. Blue coloured plot: pathways affected by genes significantly lower expressed in adult compared to L1; red coloured plots: pathways affected by genes significantly higher expressed in adult compared to L1. Only the ten most differentially expressed pathways per database are shown. RNA sequencing data analysed using ExpressAnalyst was further analysed with EnrichR using R Version 4.4.1.

### Identification of novel diagnostic candidates

The highly expressed genes in adult GW females were screened for putative signal peptide or transmembrane (TM) domains, which may indicate export or surface exposure – features that could make them potential targets of the host immune system (S1 Table). Additionally, we examined the number of orthologs across other helminths to assess the degree of conservation or uniqueness within the phylum Nematoda. We identified several genes with high expression in adult GW females and a low number of orthologs in other nematodes including DME_0000799001 (encodes for NADH dehydrogenase subunit 1) and DME_0000835601 (encodes for a transmembrane protein with unknown function) both with predicted TM domains and DME_0000533801 (encodes for TIL domain-containing protein) with a predicted signal peptide. Among currently validated diagnostic targets [43–45], domain of unknown function 148 protein (DUF148) ranked among the top ten most highly expressed genes in GW adult females. However, thioredoxin like protein 1 (TRX), another diagnostic target under investigation [45], showed low transcript expression in the adult stage.

## Discussion

Since the emergence of GW infection in dogs and other animals, there has been an increasing need to refine research tools and re-evaluate the GWEP research agenda [17]. This includes a better understanding of GW biology and the identification of novel diagnostic and therapeutic targets [17]. In alignment with these new research needs, we conducted a comparative whole transcriptome analysis of two important developmental stages of GW - adult female and L1. PCA revealed that L1 pools clustered together, indicating a high degree of similarity in their transcriptome profiles. However, the two adult female GW formed distant clusters which may reflect individual variation in gene expression or potential contamination by remaining L1 inside the female worm body. Notably, we observed that approximately 80% of genes are expressed in L1 compared to 70% in adult female, reflecting the higher metabolic activity of L1 relative to adult female.

Like all nematodes, GW progresses through five developmental stages (L1-L5 or adult). These transitions from one stage to the next require molting of the collagen-rich cuticle. We analyzed the transcriptome of two individual adult females (with 3 technical replicates for each worm) retrieved from an experimentally infected ferret along with five L1 pools isolated from these two female GW. Differential gene expression analysis and gene ontology analysis of up- and downregulated genes in adult female compared to L1 highlighted the expression of cuticle-associated genes in the adult stage. The nematode cuticle, largely composed of collagens, is critical for the maintenance of body morphology and broadly serves as interface for parasite-host or environmental interaction [46, 47]. After mating, the GW female migrates extensively through the host tissues to reach the subcutaneous space while developing and producing L1. Cuticle integrity is critical during this phase, as the worm grows up to one meter in length and remains in the tissue for several months. Additionally, it must retain millions of L1 within its body until the time of emergence. The cuticle also plays a key role in expulsion and release of L1 to the environment through interactions with the nematode muscles. Our findings suggest that the cuticle may represent a promising therapeutic target for treating GW infections by specifically interfering with the release of L1.

We further investigated genes that are highly expressed in adult GW females to identify novel diagnostic markers. The genes encoding for NADH dehydrogenase subunit I, TIL-domain containing protein and a transmembrane protein with unknown function might be exposed to the host’s immune system and are potential novel diagnostic candidates. Among the current diagnostic candidates of GW, DUF148 was highly expressed in adult female. This is consistent with prior findings from immunoproteomic studies, which showed strong reactivity with sera from dogs with a history of GW infection [44, 45]. While DUF148 is abundantly expressed at the transcript level, it has 160 orthologs in other nematodes, raising concerns about possible cross-reactivity and false positivity in diagnostic screens. Our previous assessment also confirmed that DUF148 cross-reacts with antibodies from dogs infected with *Brugia pahangi* and gastrointestinal nematodes [45]. Unlike DUF148, TRX was not highly expressed at the transcript level in this study, which may explain its poor performance as a diagnostic marker [45].

Among the genes and pathways upregulated in L1 compared to adult female GW, we observed significant enrichment in DNA replication, transcription and translation, as well as the inhibition of protein ubiquitination. Notably, within translation, the formation of the 60S and 40S ribosomal subunits, as well as translation initiation and termination, were significantly enriched. This elevated expression of genes within transcription and translation suggests that L1 are metabolically prepared for rapid transition needed for survival in the freshwater environment, successful infection of copepod IH, and subsequently molting to the next stages (L2 and L3). Similar patterns have been observed in other parasitic nematodes: for instance, in the barber’s pole worm *Haemonchus contortus,* genes associated with DNA replication and transcription are upregulated during transition from egg to L1, highlighting the developmental shift to a motile and feeding stage [48, 49]. In the filarial nematode *Brugia malayi*, distinct transcriptional profiles correspond to different lifecycle stages and transcription factors and eukaryotic translation initiation factors were highly expressed in microfilariae [50].

While this comparative transcriptomic study provides valuable insights into gene expression profile of GW at adult female and L1 stages, several limitations should be noted. Our study relied on limited biological replicates particularly for the adult female from a single mammalian host (ferret) that is not an actual reservoir host for GW in the endemic countries. Additionally, we used frozen nematode segments and L1 which might affect transcriptional profile [51]. The presence of retained L1 in adult female samples may confound stage-specific interpretation. Finally, the current *D. medinensis* genome is poorly annotated, which limits functional interpretation. Future studies integrating proteomics and spatial transcriptomics using fresh samples from actual reservoir hosts could help resolve these limitations and provide a more comprehensive view of gene function and regulation in *D. medinensis*.

GW disease remains the only human parasitic disease targeted for eradication that lacks a vaccine, drug, or widely available diagnostic test to detect prepatent infections. Our comparative transcriptome study sheds light on the biology and development of GW at two important stages, the adult female and L1. This transcriptome will serve as a valuable resource for future studies aimed at identifying drug targets and diagnostic markers.

## Supporting information

**S1 Fig. Volcano plot of differentially expressed genes in adult compared to larval L1 stages of *Dracunculus medinensis*.** Blue: significantly lower expression in adult compared to L1; red: significantly higher expression in adult compared to L1; grey no significant change in adults compared to L1. Thresholds: adjusted P-value of 0.05 and log2 fold change set to 0. Analysis and figure generation was done with ExpressAnalyst (Seq2FunM 2.0.5).

**S2 Fig. Heatmaps of Gene Set Enrichment Analysis (GSEA) using two databases.** Analysis was done with ExpressAnalyst (Seq2FunM 2.0.5).

**S3 Fig. Dot plot of Pathway enrichment analysis of differentially expressed genes in adult compared to larval L1 stages of *Dracunculus medinensis*.** Blue coloured plot: pathways affected by genes significantly lower expressed in adult compared to L1; red coloured plots: pathways affected by genes significantly higher expressed in adult compared to L1. Only the 20 most differentially expressed pathways of each database are shown. RNA sequencing data analysed using ExpressAnalyst was further analysed with EnrichR using R Version 4.4.1.

## Acknowledgements

We would like to thank Chadian GWEP that collected the parent worm used for our study. We also thank Elias Topo and Meriam N Saleh from Texas A&M University for technical support. We are grateful to the Carter Center staff for critical reading of our manuscript.

## Data availability statement

All relevant data are within the paper or supporting information files, except for the RNA-seq data that are available at NCBI GEO database under accession number (GSE302616).

## Authors’ contributions

**Conceptualization:** Hassan Hakimi, Lucienne Tritten, Guilherme G. Verocai

**Data curation:** Hassan Hakimi, Sabina Beilstein, Lucienne Tritten

**Formal analysis:** Hassan Hakimi, Sabina Beilstein, Lucienne Tritten, Guilherme G. Verocai

**Funding acquisition:** Lucienne Tritten, Guilherme G. Verocai

**Investigation:** Hassan Hakimi, Sabina Beilstein

**Methodology:** Hassan Hakimi, Michael J. Yabsley, Christopher A. Cleveland, Guilherme G. Verocai

**Project administration:** Hassan Hakimi, Guilherme G. Verocai

**Resources:** Lucienne Tritten, Guilherme G. Verocai

**Supervision:** Lucienne Tritten, Guilherme G. Verocai

**Validation:** Hassan Hakimi, Sabina Beilstein, Lucienne Tritten, Guilherme G. Verocai

**Visualization:** Hassan Hakimi, Sabina Beilstein

**Writing – original draft:** Hassan Hakimi, Sabina Beilstein

**Writing – review & editing:** Hassan Hakimi, Sabina Beilstein, Michael J. Yabsley, Christopher A. Cleveland, Guilherme G. Verocai

